# Towards Compilation of Balanced Protein Stability Datasets: Flattening the ΔΔG Curve through Systematic Under-sampling

**DOI:** 10.1101/2021.09.17.460216

**Authors:** Narod Kebabci, Ahmet Can Timucin, Emel Timucin

## Abstract

Protein stability datasets contain neutral mutations that are highly concentrated in a much narrower ΔΔG range than destabilizing and stabilizing mutations. Notwithstanding their high density, often studies analyzing stability datasets and/or predictors ignore the neutral mutations and use a binary classification scheme labeling only destabilizing and stabilizing mutations. Recognizing that highly concentrated neutral mutations would affect the quality of stability datasets, we have explored three protein stability datasets; S2648, PON-tstab and the symmetric S^sym^ that differ in size and quality. A characteristic leptokurtic shape in the ΔΔG distributions of all three datasets including the curated and symmetric ones were reported due to concentrated neutral mutations. To further investigate the impact of neutral mutations on ΔΔG predictions, we have comprehensively assessed the performance of eleven predictors on the PON-tstab dataset. Correlation and error analyses showed that all of the predictors performed the best on the neutral mutations while their performance became gradually worse as the ΔΔG of the mutations departed further from the neutral zone regardless of the direction, implying a bias towards dense mutations. To this end, after unraveling the role of concentrated neutral mutations in biases of stability datasets, we described a systematic under-sampling approach to balance the ΔΔG distributions. Before under-sampling, mutations were clustered based on their biochemical and/or structural features and then three mutations were systematically selected from every 2 kcal/mol of each cluster. Upon implementation of this approach by distinct clustering schemes, we generated five subsets varying in size and ΔΔG distributions. All subsets notably showed amelioration of not only the shape of ΔΔG distributions but also other pre-existing imbalances in the frequency distributions. We also reported differences in the performance of the predictors between the parent and under-sampled subsets due to the enrichment of previously under-represented mutations in the subsets. Altogether, this study not only elaborated the pivotal role of concentrated mutations in the dataset biases but also contemplated and realized a rational strategy to tackle this and other forms of biases. Under-sampling code is available on GitHub (https://github.com/narodkebabci/gRoR).

## INTRODUCTION

Investigating how an amino acid substitution affects protein stability is an important task refining our understanding of protein folding.^1, 2^ There are, both experimental and theoretical, ways to accurately determine the change in folding free energy due to a mutation (ΔΔG).^3–5^ Nonetheless, these approaches, either site-directed mutagenesis experiments or free energy calculation methodologies, are usually expensive, arduous and time-consuming. Thus, alternative computational approaches, also known as protein stability predictors, that both expedite the calculations and reduce the costs have been developed. ^6–8^ While these predictors implement distinct algorithms to compute ΔΔG, they usually rely on the available empirical datasets for either parametrization of their functions or training of their models. This reliance basically accentuates the significance of dataset quality for development of accurate predictors.^9^

Almost all of the mutation data utilized for development of the stability predictors came from one of the largest collections, ProTherm. ^10^ However, a number of issues have been previously raised related with the quality of this dataset and its subsets.^11–13^ While some of the trivial issues such as repetitions, mismatches and/or incorrect entries were easily solved, ^11^ others that were intrinsic to the empirical data were not. Of these intrinsic issues, higher abundance of destabilizing mutations than stabilizing mutations as reflected by an asymmetric ΔΔG distribution with a positive skew has been encountered in almost all mutation datasets.^14, 15^ This asymmetry was arguably considered critical as it has been associated with a bias towards destabilizing mutations for ΔΔG predictors.^16–22^

Other imbalances particularly in the amino acid frequencies that would affect accuracy of ΔΔG predictions also exist. Because the side chain of alanine is formed by a single carbon (C*β*), it was conveniently perceived as the best replacement amino acid. ^23^ As a result, mutant positions in the stability datasets constantly accommodated more alanine than any of the 19 amino acids.^24, 25^ Moreover, amino acid frequencies are not naturally balanced such that small aliphatic amino acids such as alanine occur more frequently in proteins than do large amino acids,^26–28^ also justifying high alanine incidences in the stability datasets. Albeit natural or universal, such imbalances are not desired in the stability datasets as they have been recognized as major sources of bias curtailing development of unbiased predictors.

From another perspective, the imbalances in the stability datasets would also skew the results of comparative assessment studies which test the performances of the stability predictors to pinpoint their strengths and weaknesses. ^22,24,29–32^ Hypothetically, the performance of an unbiased predictor may fall behind that of a biased predictor if they were evaluated on an unbalanced dataset. Thereof, we underscore the importance of compilation of a balanced stability dataset not only for resolving the over-fitting issues of the ΔΔG predictors but also for reliable assessment of their performances.

Compilation of a balanced dataset is a complex task that first requires spotting all of the imbalances and then resolving them. To date, many efforts have been dedicated to eliminate some of the well-known imbalances.^9,13,33,34^ One successful example is the generation of the S^sym^ dataset that overcame the asymmetry in the frequencies of destabilizing and stabilizing mutations.^33^ Another recent study investigated the dataset-dependence of the predictors and generated mutation-type balanced subsets from a curated dataset by a randomized under-sampling strategy. ^9^ Notwithstanding these efforts, other imbalances still exist in the stability datasets awaiting to be detected and resolved.

Mutations in the stability datasets are not evenly scattered over the ΔΔG range such that neutral mutations are usually highly concentrated in a much narrower ΔΔG range than other mutations. Despite this imbalance in the mutation densities, often the bias towards mutation-types was assessed through the use of a binary classification scheme which labels only destabilizing and stabilizing mutations. ^17, 18^ However, almost all mutation datasets contain mutations with their ΔΔG equal or close to zero. Assignment of these neutral mutations to either of the category would be fundamentally flawed and more importantly would hinder the impact of the density imbalance on dataset biases. Thus, a ternary classification addressing for the neutral type as well would more accurately group mutations than the binary mode. In this background, how the performance of the predictors on neutral mutations would compare with those on the destabilizing and stabilizing mutations needs further clarification.

Prompted by this necessity, this study addressed the role of neutral mutations (and mutation densities), in the biases of stability datasets by closely examining three stability datasets varying in size and quality and benchmarking eleven different ΔΔG predictors on an unbalanced dataset. From the dataset perspective, we have affirmed that concentrated neutral mutations led to a characteristic peaked and leptokurtic shape in the ΔΔG distributions, while from the predictor perspective all of the predictors were shown to be biased towards the mutations in the neutral range. Hence, after accentuating that concentrated neutral mutations, i.e the peaked shape of the ΔΔG distributions, as a critical imbalance for stability datasets, we described a systematic under-sampling approach to flatten the ΔΔG distributions by sampling of an equal number of mutations from a given ΔΔG range. This approach was implemented on the unbalanced dataset that was used for benchmarking generating different subsets. Aside from having more balanced ΔΔG shapes than the parent dataset, all of the under-sampled subsets also showed balanced distributions of the mutation types.

## MATERIALS AND METHODS

### Stability Datasets

Three mutation datasets, S2648, PON-tstab (S1564) and S^sym^ were retrieved from the VariBench^35^ (http://structure.bmc.lu.se/VariBench). The ΔΔG sign convention within these datasets is set to indicate stabilizing mutations with a negative sign and destabilizing mutations with a positive sign. Neutral mutations were referred to those with ΔΔG in the range of [−0.5 – 0.5]. All three datasets were investigated by means of their ΔΔG and amino acid frequency distributions.

### Curation of the PON-tstab Dataset

The downloaded PON-tstab was curated to keep the mutations (i) that were not repeated, (ii) did not have any mismatches and/or (iii) had a correct PDB information (see supplementary information for further details). Some of the repetitions had the same ΔΔG while some had different values. Because the difference in the ΔΔG of repeated mutations did not vary significantly in this dataset, we randomly eliminated half of the duplications and two thirds of the triplications. Every other problematic entry was closely examined by cross-checking its sequence and structure information. If a mismatch was identified to be related with inconsistencies in the residue numbering of PDB structures, the entry was corrected and kept, otherwise eliminated. Mutations with missing PDB IDs were kept if other entries from the same protein were spotted in the dataset. Otherwise no novel PDB ID information was introduced to the curated set. The curated dataset was available at the url: https://bit.ly/3xNg0tr.

### Under-sampling Strategy

Each mutation was summarized by a 2-letter representation by using either common 20-letter (20L) alphabet that can give rise to 380 different mutation groups or a 4-letter (4L) reduced alphabet that can give rise to 16 different groups. The reduced alphabet was generated based on the side chain biochemistry: *Ali* for amino acids with aliphatic (A, I, L, V, M, P, G), *Aro* for aromatic (F, Y, W), *Pol* for polar (S, T, C, N, Q) and *Cha* for charged sidechains (D, E, H, K, R). Next, each mutation group was further grouped into the subgroups based on their structural features of secondary structure and relative accessible surface area (ASA) which were determined by DSSP^36, 37^ and Biopython^38^ respectively. For both of the structural features three labels were generated. All helical structures such as *α*-helix, 3-10 helix and Π-helix were combined into the single label of *helix*, all *β*-configurations such as strands, bridges and sheets into the *sheet* label and all of the disordered regions such as turns and coils into the *loop* label. Similarly, three ASA labels were generated for the none-to-low exposed mutations with ASA less than 0.10, low-to-medium exposed mutations with ASA ranging between 0.10 and 0.50 and lastly for the medium-to-high exposed mutations with ASA higher than 0.50. In the under-sampling step, mutation sub-groups were ranked according to their ΔΔG and three mutations that had the maximum, median and minimum ΔΔG values of the same 2 kcal/mol interval were selected for every sub-group. The code is available on GitHub (https://github.com/narodkebabci/gRoR).

### Protein Stability Predictors

11 different stability predictors, namely DeepDDG,^39^ mCSM,^40^ INPS-3D, ^17^ I-Mutant 2.0,^41^ I-Mutant 3.0,^20^ SDM,^19^ MAESTRO,^42^ PoPMuSiC, ^43^ DUET,^44^ iStable^45^ and iDeepDDG,^39^ were tested on the curated PON-tstab dataset. I-Mutant 2.0 and I-Mutant 3.0, protein sequences were used as input, while for the rest of the predictors 3D structures were utilized. Predictions by I-Mutant 2.0 and I-Mutant 3.0 were done at the temperature of 25°C and pH of 7.0. Unless otherwise stated, default parameters were applied for all predictions.

### Performance Assessment

Scoring performance was assessed by the Pearson correlation coefficient (R) (Eq. 1) and the errors of the predictors were estimated by the metrics of mean absolute error (MAE) (Eq. 2) and mean signed error (MSE) (Eq. 3).

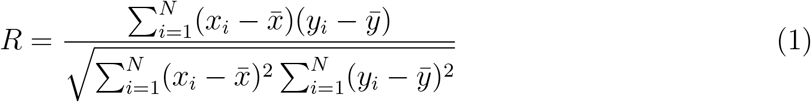

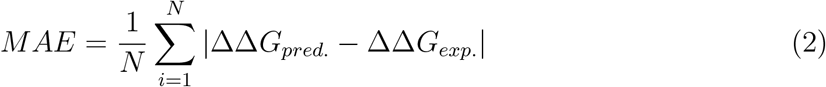

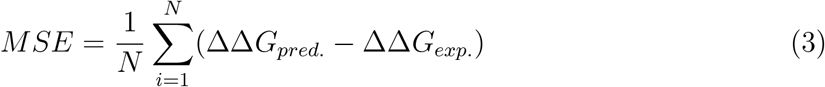

To quantify the agreement between the experimental and the computational methods, enhanced Bland-Altman plots were used to illustrate the mean difference between the experimental and predicted scores.

## RESULTS AND DISCUSSION

### Exploring a sample of stability datasets

Three datasets namely S2648, PON-tstab and S^sym^ that differed in size (Table 1) and curation status were extracted from the VariBench. ^35^ The largest of all, S2648, was a non-curated dataset^46^ while the medium-sized PON-tstab dataset was reported to be free of large redundancies^11^ and the smallest S^sym^ was purposefully built to overcome mutation-type imbalance.^33^ Exploring these datasets that comprise a decent sample of the stability data can help us pinpoint the common biases in stability datasets.

**Table 1:** Statistics of ΔΔG distributions.

| Datasets | n | $n_P$ | Median | Min. | Max. | Skewness | Kurtosis |
| --- | --- | --- | --- | --- | --- | --- | --- |
| S2648 | 2648 | 131 | 0.82 | -5.0 | 6.8 | 0.148 | 0.749 |
| PON-tstab | 1564 | 99 | 0.70 | -17.4 | 8.0 | 1.153 | 6.951 |
| $S^{\text{sym}}$ | 684 | 15 | 0 | -7.5 | 7.5 | 0 | 1.599 |
| $n_P$ is the number of proteins; median, min. and max. are in kcal/mol. | | | | | | | |

We have visualized the ΔΔG and frequency distributions of the datasets to evaluate whether they harbor any mutation-type imbalances or not (Fig. 1). Particularly, the shape of the ΔΔG distributions suggested that all datasets except for the S^sym^ were skewed towards destabilizing mutations. The skewness statistics of the distributions quantitatively indicated the same result such that the medium-sized PON-tstab had the most skewed ΔΔG distribution while the S^sym^ had zero skewness (Table 1). This imbalance as probed by the skewness statistic was recognized as the leading cause of the prediction bias towards destabilizing mutations.^16–18^

**Figure 1:**
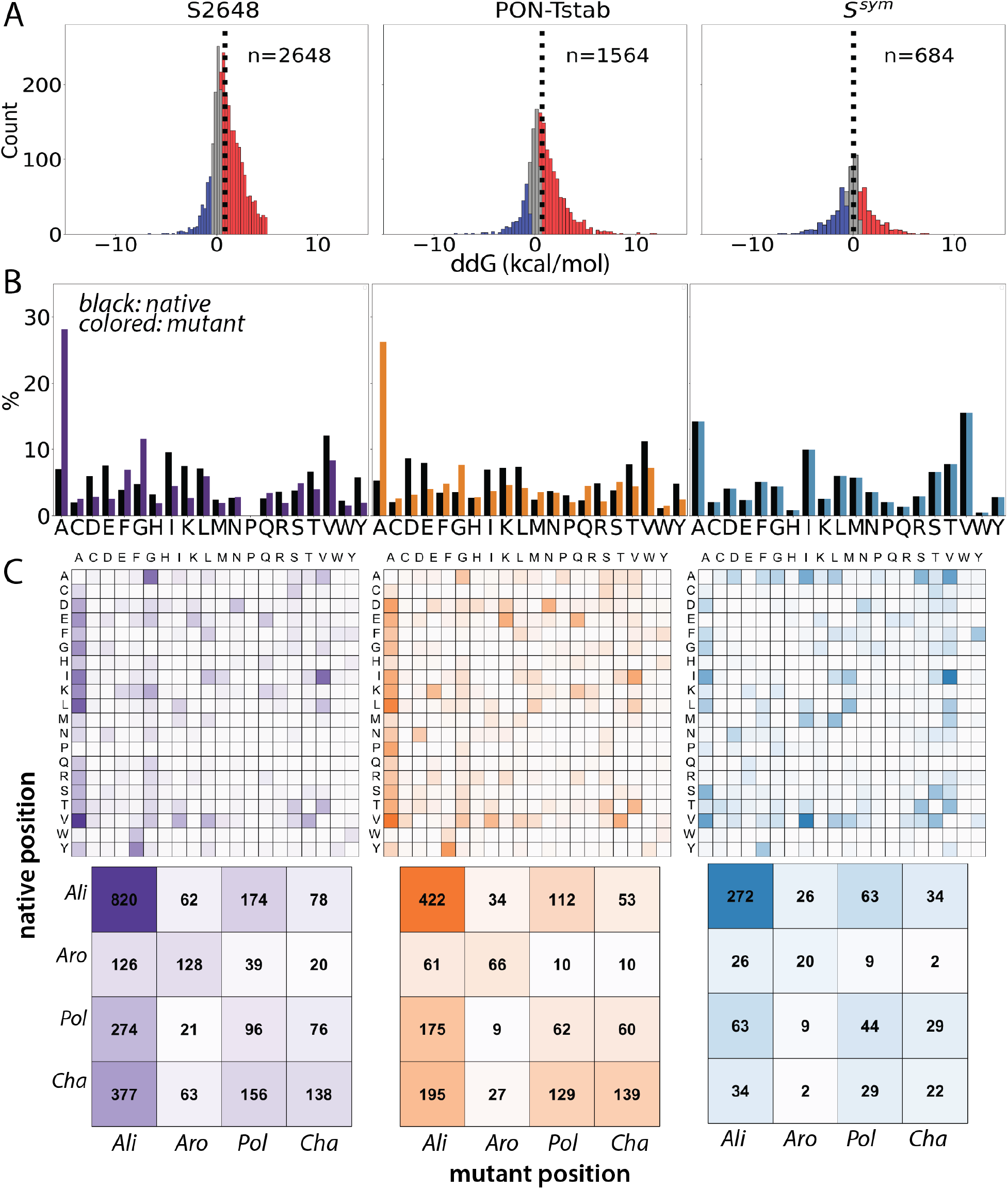
**(A)** ΔΔG distributions of three datasets; blue, red and grey colors indicate stabilizing, destabilizing and neutral mutations, respectively. Dotted lines show the median of the distributions. **(B)** Relative frequencies of amino acid in the native (black) and mutant (colored) positions. **(C)** Frequencies of the mutation groups formed by (top) 20L and (bottom) 4L alphabet.

According to amino acid frequencies, none of the datasets including the S^sym^ showed balanced frequencies (Fig. 1B). While alanine and valine were highly abundant in all datasets, other amino acids such as histidine, proline and tryptophan were scarce. Especially, the largest dataset S2648 did not contain any proline at all. Despite the *de-facto* perception of alanine as the best replacement amino acid, histidine and asparagine have been shown to be better than alanine to study mutation effects. ^47^ However, these amino acids were much less frequently sampled than alanine for all datasets. Furthermore, the S2648 and PON-tstab sampled significantly higher alanine in the mutant position than they did in the native position while the S^sym^ predictably showed symmetric frequencies across native and mutant positions for all amino acids.

To further inspect the distributions of individual mutations, we encoded the substitutions by using the common 20L and produced checkerboard plots based on the absolute frequencies (Fig. 1C-top). The most abundant substitutions were VA and IV for all datasets while more than half of the mutation types were not sampled at all by any of the datasets. Nevertheless, expecting a balanced distribution of the mutation types based on 20L alphabet is highly unrealistic for any dataset. Thus, we accordingly used a 4L reduced alphabet based on the side-chain biochemistry that could form 16 different mutation groups (Fig. 1C-bottom) for a more realistic assessment of the mutation-types.^9^ All three datasets showed deviated frequencies from the balanced dataset (Fig. S1). Notwithstanding its symmetric shape and smallest size, the S^sym^, among other two datasets, had the most deviated frequencies (Fig. 1C-bottom). Approximately 40% of the S^sym^ dataset (272/684) were aliphatic-to-aliphatic conversions while this ratio was reduced to 30% in the S2648, 20% in the PON-tstab and 11% in the balanced dataset. Furthermore, charged-to-aromatic conversions that comprised to only 0.3% of the S^sym^ accounted for higher portions of the other two datasets (0.8%) and the balanced one (3.9%). High frequencies of alanine, valine and isoleucine in both of the native and mutant positions of the S^sym^ might explain its larger divergence from the balanced dataset than the other two datasets, because these amino acids were abundant only in the mutant positions of the S2648 and PON-tstab (Fig. 1B).

We also analyzed the scoring performance based on the protein types (Fig. S3). All predictors consistently showed good performance on the proteins with high number of mutations such as lysozyme and barnase while the scoring performance greatly varied and became poorer on the proteins that were represented with a few number of mutations. This trend, the gradual decrease in the scoring performances with decreasing number of the mutations, was likely to be due to the differences in the mutation-type ratios of different proteins, the observation which was another indication of the bias toward mutation-types. Overall we confirmed the presence of the well-known mutation-type imbalances in all three datasets including the symmetric and curated datasets. Apparently, compilation of a balanced dataset requires attention to a list of features,^48^ otherwise ensuring a symmetric distribution and/or removing of large redundancies does not necessarily solve all of the quality issues.

### Peaked ΔΔG distributions as another source of bias

While some predictors tend to label neutral mutations^11,20,24,49^ often a binary classification is used to group mutations, i.e. destabilizing/stabilizing.^19,39–41^ However, almost all stability datasets contain mutations with their ΔΔG close or equal to zero reflecting the necessity of an exclusive label for neutral mutations. For instance, approximately 1-to-2% of mutations analyzed here had zero effect on the stability (Fig. S2). Although these mutations were generally considered as above zero and labeled as destabilizing mutations, we stress that assignment of these zeros to either of the mutation category is not correct. Accordingly, we used an additional label for the neutral mutations that had ΔΔG values between −0.5-0.5 kcal/mol. ^49^ While alternative ranges can be used to delimit neutral mutations,^9, 50^ we underscore the ultimate importance of an additional label for the neutral mutations and stress the insufficiency of the dichotomous mutation category for analyzing dataset biases.

We found 726, 468 and 235 neutral mutations in the S2648, PON-tstab and S^sym^ respectively. Although these neutrals corresponded to approximately one third of the datasets, at first implying no particular imbalance in the frequencies (Fig. S2), they were concentrated in a much narrow ΔΔG range than the other two mutation types for all datasets. Explicitly, for the S2648 dataset, destabilizing mutations, corresponding to 60% of the dataset, were distributed over the range of 6.29 kcal/mol, while stabilizing mutations, accounting for 15% of the S2648, spanned a similar range of 4.49 kcal/mol. As a result, the concentration of neutral mutations were noted to be much higher than those of the destabilizing and stabilizing mutations in the S2648 dataset. More extreme observations were made for other datasets confirming that all three datasets had highly concentrated neutral mutations. The ΔΔG distributions with highly concentrated neutral mutations would over-represent neutral and mildly stabilizing/destabilizing mutations while they would systematically under-represent highly stabilizing/destabilizing mutations. Thereof, in line with the desired characteristics of the frequency distributions,^9^ one would expect from a balanced dataset to have a less peaked ΔΔG distributions with less neutral mutations.

Similar to the use of skewness as the measure of symmetry, the kurtosis statistic of ΔΔG distributions could be used to quantitatively evaluate the peaked-ness of the distributions and thus the concentrations of neutral mutations. As the concentration of neutral mutations increases, the kurtosis (peaked-ness) of the ΔΔG distributions increases. Thus, we have calculated the kurtosis excess values for all distributions (Table 1) which produced parallel results with the visual assessments (Fig. 1A). All three datasets exhibited positive kurtosis, particularly the distribution of the PON-tstab had the highest excess kurtosis resulting in the most leptokurtic shape while the S2648 had the best shape among all three with the lowest excess kurtosis. Theoretically, a platykurtic ΔΔG distribution with a negative kurtosis provides a more balanced representation of the mutation types than does the mesokurtic or leptokurtic distributions with a zero or positive excess kurtosis. Therefore, we confirmed that none of the datasets showed a balanced ΔΔG distribution, i.e. negative kurtosis.

In other words, the ΔΔG distributions with positive kurtosis would over-represent neutral and mildly stabilizing/destabilizing mutations while they would systematically underrepresent highly stabilizing/destabilizing mutations. Nevertheless, these under-represented mutations are of high interest particularly for the disease-related cases as extreme ΔΔG changes could lead to severe pathological consequences by severely affecting the structure, i.e. highly destabilizing or stabilizing effect. ^51, 52^ In fact, a medium level correlation was established between the probability of being a pathogenic variant and having a large perturbation to the protein stability. ^53^ Thus, the stability datasets with a uniform ΔΔG distribution, i.e. platykurtic, would better represent distinct mutation categories such as those with extremely high or low ΔΔG values. Overall, these and other stability datasets in the VariBench ^35^ showing leptokurtic ΔΔG distributions are carriers of a possible bias to over-represent neutral ΔΔG range.

### Benchmarking the stability predictors on the curated PON-tstab

To assess the performance of 11 ΔΔG predictors, we have recruited the PON-tstab dataset that had the most leptokurtic ΔΔG shape. Before this, the original PON-tstab was closely inspected and some inconsistencies were spotted (Table S1). Essentially, 113 mutations were removed from the dataset due to three main issues; repetition, mismatch or PDB related, resulting in a total of 1451 mutations from 89 proteins. Overall ΔΔG and frequency distributions of the curated PON-tstab were highly similar to the original dataset (Tables 1 and 2) (See supplementary information further details of the curation and the curated list of mutations).

**Table 2:** Characteristics of ΔΔG distributions of the under-sampled datasets generated in this study.

| Dataset | n | n<sub>P</sub> | Median | Skewness | Kurtosis | Skewness* | Kurtosis* |
| --- | --- | --- | --- | --- | --- | --- | --- |
| cur. PON-tstab | 1451 | 89 | 0.70 | 1.187 | 6.931 | 0.825 | 3.825 |
| 20L | 910 | 68 | 0.78 | 1.025 | 4.964 | 0.682 | 2.508 |
| 4L | 186 | 49 | 0.75 | 0.822 | 2.917 | 0.335 | 0.786 |
| 4L/SS | 409 | 61 | 1.19 | 0.756 | 3.380 | 0.379 | 1.415 |
| 4L/ASA | 385 | 70 | 1.10 | 0.768 | 3.027 | 0.411 | 1.200 |
| 4L/SS/ASA | 699 | 80 | 1.00 | 0.938 | 4.242 | 0.587 | 1.946 |

All datasets have the same range of -8.0-17.40.
\*Re-calculated skewness and kurtosis values after removal of the extreme value of 17.40.

Pearson’s correlation coefficient (*R*) is probably the most widely used metric for performance evaluation of the predictors.^54^ Based on the assessment by this metric (Fig. 2A), we reported the I-Mutant2.0 as the best performer. Notwithstanding sharing similarities in their prediction algorithms, the performance of I-Mutant2.0^41^ and I-Mutant3.0^20^ noticeably differed from each other such that I-Mutant3.0 had a much lower scoring performance (*R* = 0.58) than I-Mutant2.0 (*R* = 0.74) (Fig. 2A). I-Mutant2.0 has been followed by iStable while SDM was the worst performer. SDM’s performance was slightly surpassed by INPS and Maestro. Scoring performance of the rest of the predictors remained in the *R* range of 0.50 − 0.64. Among these mediocre performers, two young predictors, ^39^ DeepDDG, a neural network model, and iDeepDDG, a meta-predictor using the outputs of mCSM, SDM, and DUET, had comparable scoring performances. DUET, another meta-predictor, combining SDM and mCSM scores showed a slightly lower performance than iDeepDDG, reflecting the subtle advantage of iDeepDDG over DUET on this dataset.

**Figure 2:**
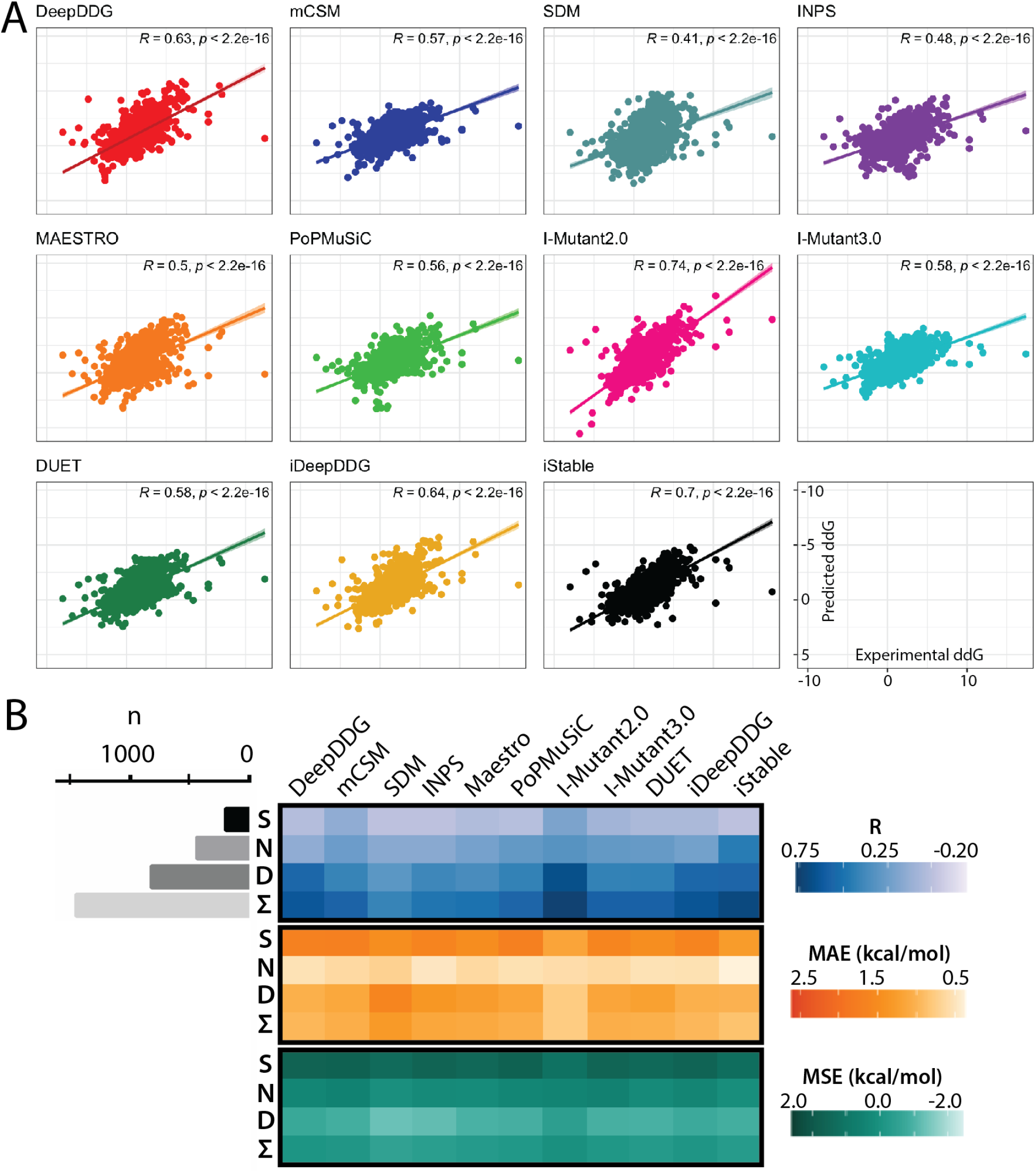
**(A)** Scoring performance of 11 predictors on the curated PON-tstab by Pearson’s R. **(B)** Scoring performance (R) and error (MAE and MSE) analyses based on mutation types (S: stabilizing, D: destabilizing, N: Neutral and Σ: Total). Number of mutations were illustrated as bar plot on the left.

To quantify the prediction errors, we have computed MAE and MSE for the entire dataset (Fig. 2B) which were in close agreement with the correlation analysis (Fig. 2A) and results of previous benchmarks.^9, 39^ I-Mutant2.0 had the lowest error of 0.77 kcal/mol while SDM had the highest error (1.31 kcal/mol), marking the lower and upper MAE bounds of this benchmark. Similarly, I-Mutant2.0 had the lowest and the single positive MSE (0.05 kcal/mol) while SDM had the largest MSE of −0.53 kcal/mol. Overall, MSE analysis implied that all of the tools showed an overall balanced prediction for the entire dataset.

To examine how the performance of the predictors varied based on mutation types, we computed the correlations for three mutation types; destabilizing, neutral and stabilizing (Fig. 2B). Almost all predictors showed an apparent preference, higher correlations, towards destabilizing mutations followed by neutral mutations while the correlations for the stabilizing mutations were the weakest for all predictors (Fig. 2B). Although this trend in the correlations was previously noted,^17, 18^ error analyses contradicted the correlation results as such the neutral mutations which had the second best scoring performance (R) were predicted with the smallest error (Fig. 2B). Despite this contradiction, stabilizing mutations were again the most problematic predictions that consistently produced the weakest correlations and largest absolute errors. Nonetheless, a particular trend in the MSE heat-map were reported that all predictors consistently made a negative signed error for the destabilizing mutations but a positive one for the stabilizing mutations reflecting that the predictors under-estimated destabilizing mutations and over-estimated stabilizing mutations. In other words, predictors tended to produce a ΔΔG score closer to zero regardless of the sign of the mutation. To illustrate the numerical agreement between the scores, we have used Bland-Altman (BA) plots (Fig. S4). While the bias of BA analysis (mean difference) was in fact the same as the MSE,^56, 57^ these plots were used to visualize the mutations falling out of the agreement limits (LOA). Particularly, highly destabilizing and stabilizing mutations with extreme ΔΔG changes resided outside the LOA whilst the neutral and mildly destabilizing/stabilizing mutations were predicted within the LOA.

Given the trend detected in the MSE and BA plots, we computed the correlations and errors for every 1 kcal/mol interval (Fig. 3). Contrary to our initial observations (Fig. 2B) and previous observations,^17, 18^ the mutations in or close to the neutral range produced the strongest correlations for every predictor. Otherwise, destabilizing mutations with ΔΔG higher than 2.5 kcal/mol showed no, if not negative, correlations at all. Similarly, we observed a heterogeneity in the correlations of the stabilizing mutations such that the 7 of 11 predictors produced a medium level correlation for the stabilizing mutations close to the neutral range (−1.5 – −0.5) while none of the predictors showed coherent performances towards for highly stabilizing mutations.

**Figure 3:**
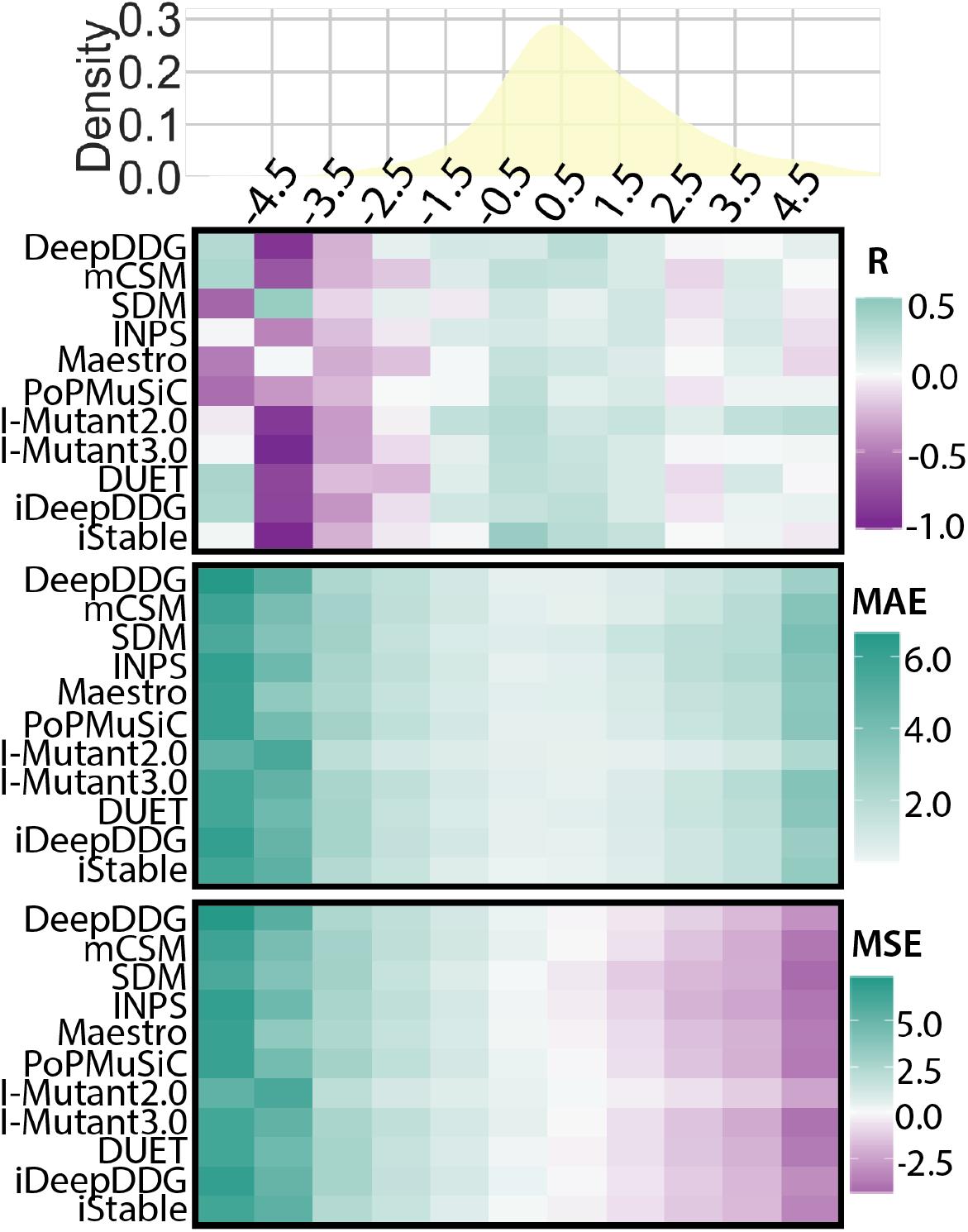
Scoring performance (R) and error (MAE and MSE) assessment based on ΔΔG. Each heatmap column was aligned to the KDE plot^55^ to indicate the mutation density for the given ΔΔG interval.

MAE analysis further corroborated that as the experimental ΔΔG of a mutation departed from zero regardless of its sign, the error associated with its ΔΔG prediction systematically inflated for every predictor. While MSE heat-map confirmed that every predictor over-estimated stabilizing mutations and under-estimated the destabilizing mutations. According to the correlation and error heat-maps computed for every 1 kcal/mol interval, all predictors made smaller errors as the density of the mutations increased. Overall, we underscore the need for the recapitulation of the well-recognized prediction bias towards destabilizing mutations, at least for this benchmark, as the bias towards concentrated mutations such as neutral and mildly destabilizing/stabilizing mutations.

### Systematic under-sampling of PON-tstab leads to subsets with more balanced ΔΔG distributions

Among three datasets, the medium-sized PON-tstab had the most peaked and skewed ΔΔG distribution (Table 1 and Fig. 1) along with a non-uniform amino acid distribution which would least align with the desired characteristics of a balanced dataset. Thus, we recruited the curated PON-tstab as the toy dataset to test our under-sampling strategy (Fig. 4).

**Figure 4:**
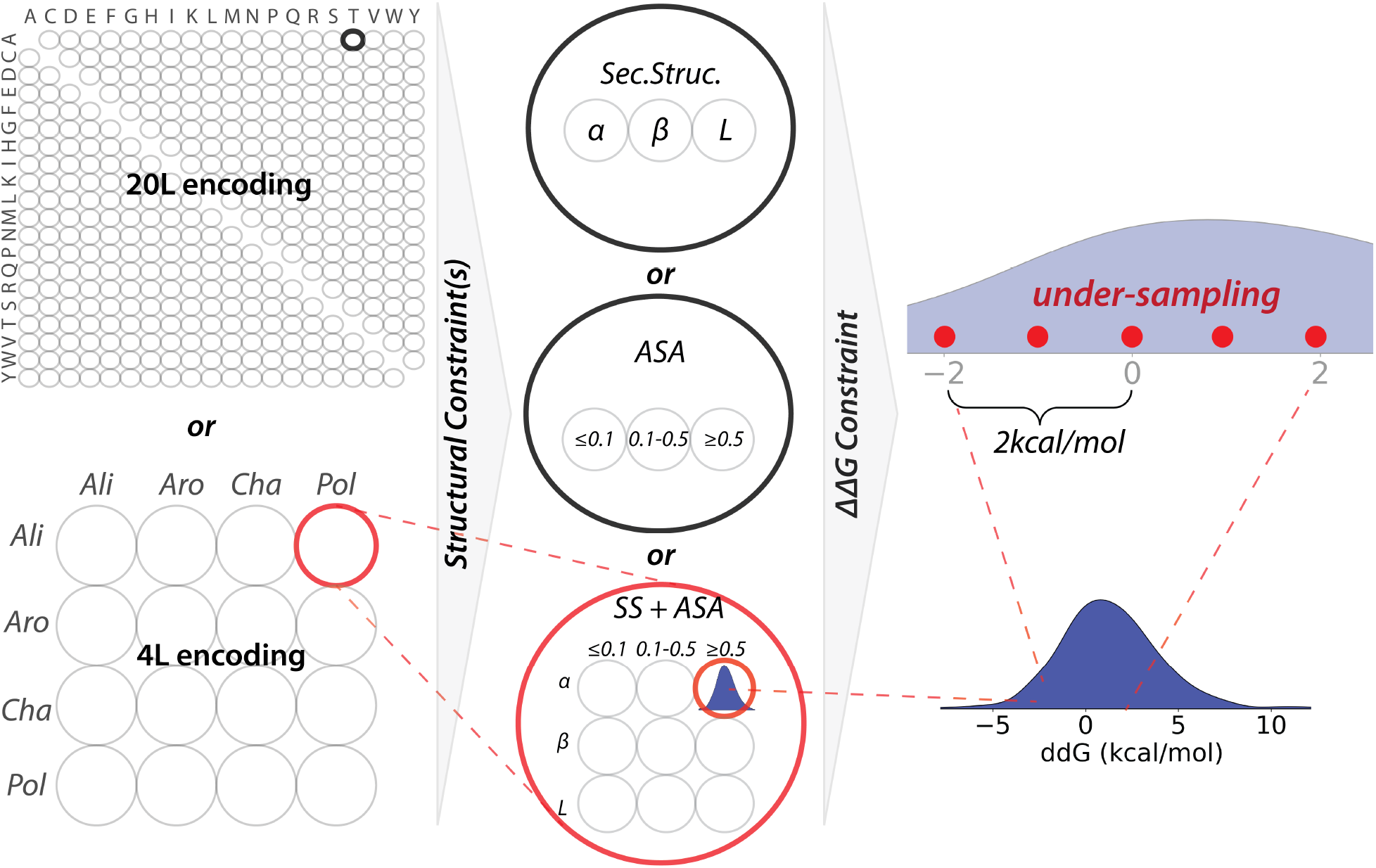
Systematic under-sampling workflow. (Left) Either 20L or 4L alphabet was used to group mutations. (Center) Structural constraints were used to further group the mutation groups, forming subgroups based secondary structure and/or ASA similarities. (Right) Stability constraint was applied to rank the mutations in every subgroup. From every 2 kcal/mol, three mutations with the maximum, median and minimum ΔΔG were selected.

Before under-sampling, we have clustered the mutations based on their biochemical and/or structural properties. To achieve the former, a reduced alphabet of size of four was used.^9, 58^ Despite availability of alternative reduced amino acid alphabets, ^59, 60^ we have chosen the small 4L alphabet based on the side chain biochemistry which effectively reduced the number of mutation groups and still accounted for biochemical similarity within the mutation group (Fig. S1). For the latter, two structural features; secondary structure and relative ASA which were illustrated on 3 example proteins; barnase, nuclease, PSBD (Fig. S5) were utilized. By combination of two alphabets and two structural constraints, we have generated five distinct subsets from the curated PON-tstab (Fig. 5). First, we have generated the 20L subset by under-sampling of every 2 kcal/mol of all mutation groups formed by 20L.

**Figure 5:**
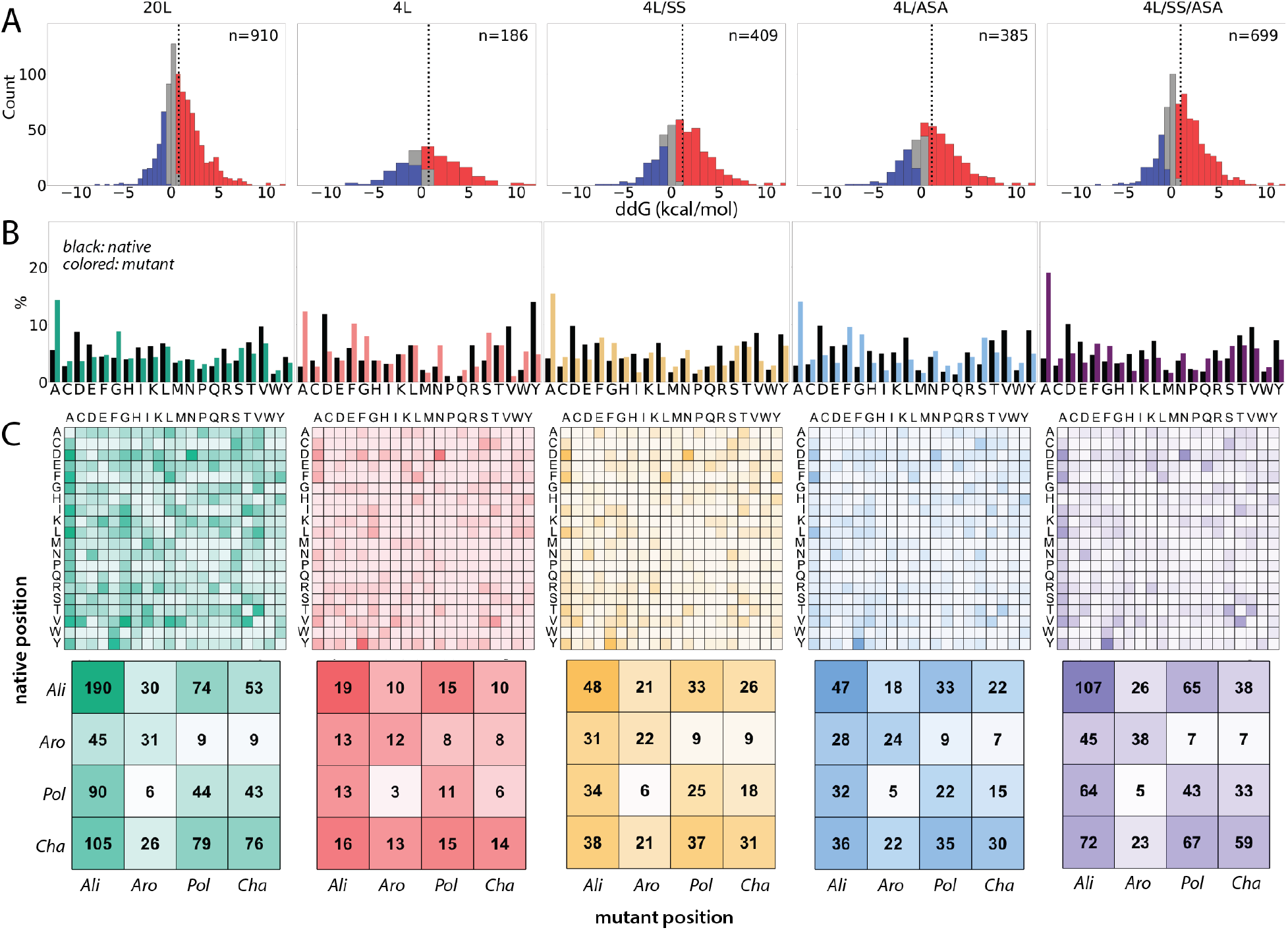
Five different under-sampled datasets. Vertical panels representing different undersampling trials show **(A)** ΔΔG distributions, **(B)** relative amino acid frequencies, **(C)** absolute frequencies according to (top) 20L and (bottom) 4L alphabets.

In the curated PON-tstab we have spotted that 181 of the possible 380 mutation types were either missing or sampled once. Thus, we did not apply any structural constraints to the 20L mutation groups which would further reduce the data points in the groups. For the second subset, we have used 4L alphabet to form 16 mutation groups which notably increased the number of mutations in each group. Upon sampling of 3 mutations from every 2 kcal/mol of each of the 16 mutation group, we have compiled the smallest subset of 4L. Owing to the small size of the reduced alphabet, we have observed that all of the possible 16 groups were sampled by the curated PON-tstab. The third and fourth panels in Fig. 5 showed the subsets constructed by applying one of the structural constraints to the 4L subset, while the fifth and last subset was formed by applying both of the structural constraints to the 4L subset. Within this 4L/SS/ASA subset, approximately half of the mutations were eliminated yet the under-sampled dataset conserved the same ΔΔG range (Table 2). Visually, we showed the sampled and eliminated mutations on the three example proteins for the same subset (Fig. S5). Particularly, the NMR structure of PSBD displayed that only a few mutations were selected from its terminal helix which contained highly similar substitutions mutations according to their biochemical and structural features.

Inspection of the ΔΔG distributions of the under-sampled subsets suggested amelioration of almost all of the statistics of the PON-tstab (Table 2). Particularly, the skewness and kurtosis measures were reduced in every subset regardless of the applied under-sampling scheme. Frequency distributions of amino acids were also improved such that compared to the parent dataset (Fig. 1B-C), under-sampled subsets had more balanced frequencies across mutation groups based on both alphabets (Fig. 5B-C). Although among all 5 subsets, the subset 4L was the most balanced subset with the highest reduction in the kurtosis and displayed the most improved statistics (Table 2), it had the smallest size which was reduced to only ~10% of the parent dataset (Table 2). Because larger datasets that ensure higher statistical power than smaller datasets would be more preferable for training ΔΔG predictors,^61^ we particularly considered the subsets of 4L/SS, 4L/ASA or 4L/SS/ASA as the optimal under-sampled subsets of the PON-tstab.

Our strategy evenly sampled 3 data points from every 2 kcal/mol interval of the mutation groups formed based on biochemical and/or structural similarities. Thus, consistently denser mutations were eliminated more than less dense mutations. Due to the highest concentration of neutrals in the parent dataset, highest number of eliminations were made from the neutral mutations, leading to a discernible reduction in the kurtosis of ΔΔG distributions (Table 2). However, although much less than the neutral mutations, some destabilizing and stabilizing mutations were also eliminated, leading to only a slight improvement in the skewness, the ratio of destabilizing-to-stabilizing and even a decline in the central tendency measures as the median and mean of the subsets slightly shifted towards destabilizing range (Table 2, Figs. 5 and S2). Although we did not implement here, some measures might be undertaken to strictly balance the destabilizing and stabilizing mutations. For instance, alternative subsets can be formed without making any eliminations from stabilizing mutations or stabilizing mutations can be selected from a narrower interval than 2 kcal/mol. Furthermore, the PON-tstab had an asymmetrical ΔΔG range of −8 – 17.4 kcal/mol and thus our under-sampling strategy led to collection of more destabilizing mutations than stabilizing mutations. However if the ΔΔG range of the parent dataset was symmetrical, eliminations from destabilizing and stabilizing mutations would be more balanced. While these maneuvers would have better improved the skewness and central tendency measures of the subsets, we also noted that elimination of the extreme values or outliers would contribute too. As such, removal of the extreme value, 17.4 from the destabilizing mutations, greatly optimized the shape of the distributions by further reducing the skewness (Table 2).

Having observed that all predictors systematically made smaller errors towards neutral mutations (Fig. 3), we linked this bias to the leptokurtic shape of the ΔΔG distributions (Fig. 1). Hence, accordingly we pondered an under-sampling approach to construct subsets with diluted neutral mutations, i.e. less peaked ΔΔG distributions. While a more straight-forward approach would have served this purpose such as only selecting an even number of mutations from every 2 kcal/mol for the entire ΔΔG range, we have bothered an additional step to group mutations based on biochemical and/or structural similarities. The need for this extra step was in fact to establish a more balanced distributions based on biochemical and structural features in the resulting subsets. Considering that the parent dataset had high abundance of the mutations that were located in helical structures or core regions (Fig. S6), otherwise a crude under-sampling approach, albeit systematic, may have transferred these imbalances to the subsets. The benefit of using biochemical and structural restraints was illustrated in Fig. S6. Comparison of the subsets with the parent dataset confirmed that the under-sampling without application of any structural constraints led to 20L and 4L subsets that highly reflected the parent dataset regarding secondary structure or ASA distributions. While application of either or both of the structural constraints to build 4L/SS, 4L/ASA and 4L/SS/ASA subsets showed a more even distribution of these features.

### Attenuated prediction performances towards under-sampled subsets with flattened ΔΔG curves

We have calculated the scoring performance and errors for the under-sampled datasets (Fig. 6). The smallest subset 4L predictably showed the lowest correlations and highest absolute errors for all predictors (Fig. 6). As the size of the subsets were increased, we observed a gradual increase in the correlations and gradual decrease in the errors to reflect the values reached by the parent dataset. Although MSE analysis did not imply a notable change for the subsets compared with the parent dataset, inspection of the MSE based on mutation-types showed that under-sampling enriched the stabilizing and destabilizing mutations that were *hard-to-predict* by eliminating *easy-to-predict* ones. Particularly, both errors were reached to the highest levels for destabilizing and stabilizing mutations in the 4L subset (Fig. S7) suggesting the highest enrichment for this subset. Agreement of the predicted scores with the experimental values was also analyzed by BA plots which similarly suggested that all of the predictions for the under-sampled datasets were less precise with a larger fraction of mutations residing outside the LOAs compared with those of parent dataset (Fig. S4). Lastly, the remarkable performance of I-Mutant2.0 which was only subtly decreased in the subsets as opposed to other predictors was noted (Fig. 6), a puzzling observation that lead us to consider a probable overlap between the PON-tstab and the training dataset of I-Mutant2.0. ^9, 41^

**Figure 6:**
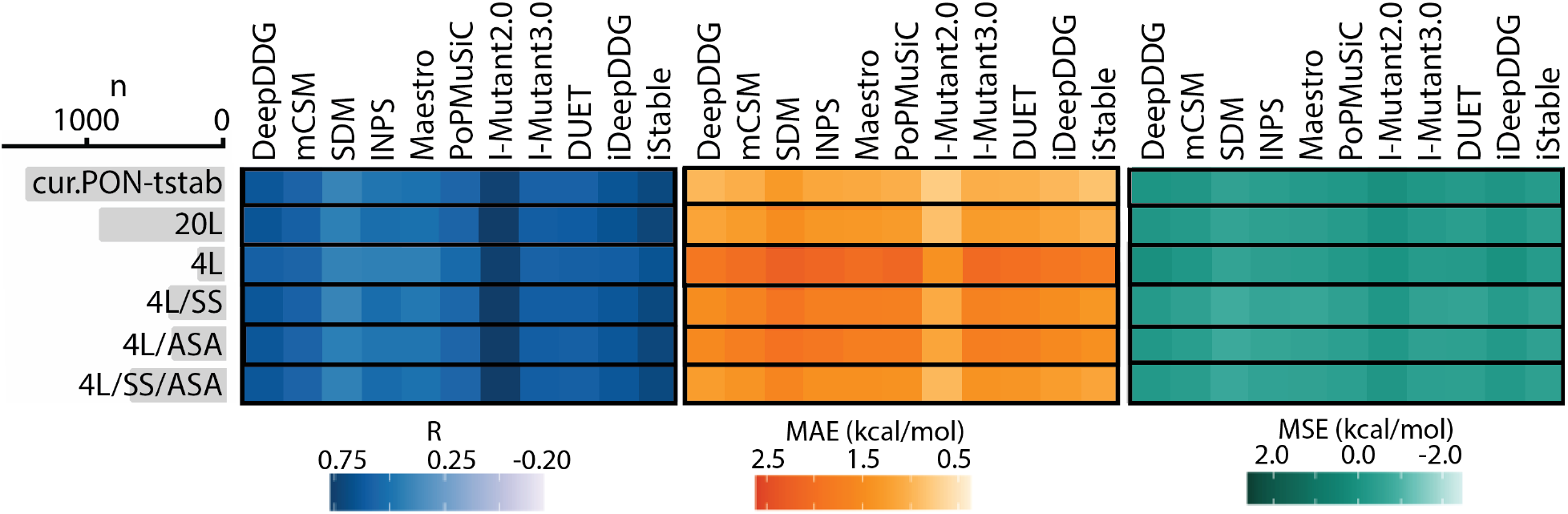
Performance analysis of 11 predictors on the curated PON-tstab and 5 under-sampled subsets by means of Pearson correlation coefficient (R), Mean Absolute Error (MAE) and Mean Sign Error (MSE).

Unlike to random under-sampling approaches,^9^ we have systematically eliminated certain mutations. The systematic nature of this approach is particularly advantageous over random under-sampling approaches because one-time implementation would be sufficient here while for random approaches, more than a few subsets need to be compiled and analyzed. In fact, we have implemented the described workflow twice for all five subsets and obtained almost the exact same set of mutations (~99%) from two independent implementations. On the other hand, because systematic elimination of certain mutations would reduce the impact of their experimental studies, it is not desired especially considering the resources donated to those studies.^48^ Nonetheless, elimination of highly similar mutations based on biochemical and structural features, as done by our approach (Fig. S6), would less likely to affect the mutation variation in the parent datasets. Furthermore, automated and flexible implementation of this approach ensure one to easily adapt it to tailor the stability subset of need. One with a much larger parent datasets such as ThermoMutDB,^25^ FireProtDB^62^ and ProtaBank^63^ can apply a more rigorous constraint selection than ours with the toy dataset PON-tstab. Different reduced alphabets can be implemented and further the sampling ΔΔG interval can be adjusted such that attenuation of this value would enforce a more stringent elimination within mutation groups and thus result in a larger subset.

## CONCLUSION

One way to overcome bias in stability datasets is to rationally design experiments that would yield data with desired characteristics such as the construction of the S^sym^. Nonetheless, many efforts have already been dedicated for compilation and curation of stability datasets. Thus, besides constructing novel datasets, rational utilization of the available datasets is a reasonable option, as attempted by this study. We stress that elaboration of the role of neutral mutations on the biases of stability datasets is the most important contribution of this study. Evidently, not only the asymmetry (skewness) but also peaked-ness (kurtosis) of the ΔΔG distributions needs specific attention to compile ideal datasets. Thus, as a tribute to the comprehensive analysis of the stability datasets by means of their distributions, we contemplated a *biasedness* score as a global measure of the dataset quality that could be computed from the statistical characteristics of datasets.

## Supporting Information Available

Supplementary information contains 1 table and 7 supplementary figures.

## Notes

https://github.com/narodkebabci/gRoR

https://bit.ly/3xNg0tr

### Competing Interest Statement

The authors have declared no competing interest.

