## Supplementary material for "Towards Compilation of Balanced Protein Stability Datasets: Flattening the ΔΔG Curve through Systematic Under-sampling": SI

#### 1 Curation of the PON-tstab

The medium-sized PON-tstab represented a curated subset of ProTherm, nevertheless, we have spotted some inconsistencies that prompted us to curate this original PON-tstab which was composed of 1564 mutations from 99 different proteins (Table S1). Overall, 269 mutations were associated with at least one problem and 113 of them were eliminated due to 3 main issues; repetition, mismatch or PDB related. Of eliminated mutations, 125 of them were repeated such that 116 mutations were duplicated and 9 were triplicated. Half of the duplication (58) and two thirds of the triplications (6) were randomly eliminated. 50 of the mismatches were corrected as they were derived from inconsistencies in the residue numbering of PDB structures. The remaining 14 with unresolved mismatches were removed. Among 145 mutations with PDB-related issues, 131 of them had their PDB IDs missing in the original PON-tstab. 48 of the IDs were corrected by cross-checking other entries of the same protein while the remaining 83 of them were eliminated. 14 mutations were eliminated due to the low PDB quality of the mutated positions. Specifically, 12 mutations from the lambda repressor whose provided PDB structure (PDB ID: 1RLP) only contained C$\alpha$ atoms and two mutations from the tryptophan synthase alpha-subunit whose structure has a missing region for the mutations (PDB ID: 1WQ5) were eliminated due to low PDB quality. This curation reduced the mutation number in the original PON-tstab to 1451 mutations from 89 different proteins (Table S1) which can be downloaded from the url:`https://bit.ly/3xNg0tr`.

Given the accumulation of structure-based predictors, only mutations with a valid PDB information were considered for our curated dataset. 50 of these eliminated mutations due to a missing PDB ID were originating from the proteins; interleukin 6, subtilisin, TEM beta-lactamase and anthranilate isomerase that were represented only by a few number of mutations in the dataset. On the other hand, 33 of the eliminated mutations in this group were obtained from the cold shock protein. Although other mutation entries for this protein are present, their PDB information (1CSP or 1C9O) does not match with any of these 33 mutations. These mutations could only be distinguished from other Csp-B entries only due to their non-standard sequence identifiers while they were listed by the same protein name. Notably, the eliminated 113 mutations were obtained from 23 different proteins while only 5 of these proteins accounted for almost half of the eliminations (49). These 5 proteins were similarly listed by a non-standard sequence identifier which in turn hampered our efforts of curation. Therefore, we highlight the importance of compilation of not only correct stability information but also correct sequence and structure information. Otherwise, correction of the missing or incorrect information in an already formed dataset would be impractical particularly for a large number of cases. Taken together with non-uniform distributions and inconsistencies that can be spotted even in the symmetrical or curated datasets, we stress that compilation of superior datasets is still a necessity.

#### 2 Scoring performance based on protein types

The proteins represented with high number of mutations closely reflected the PON-tstab dataset (Fig. 1A) and thus they showed similar, medium-to-high, level correlations for all predictors. Explicitly, the lysozyme mutations had a very similar destabilizing:neutral:stabilizing ratio (55% : 33% : 13%) to the entire PON-tstab dataset (57% : 30% : 13%). On the contrary, the proteins that were represented by only a few number of mutations failed to reflect such similar ratios. Even some proteins that had only a few mutations could randomly possess only stabilizing mutations. Two examples are apoflavodoxin and alphaparvalbumin proteins which were represented by 6 and 3 data points respectively had all stabilizing mutations. Predictors worked very poorly on these proteins, producing negatively correlated scores. Hence, we surmised that differential distributions of the mutation types in the individual proteins led to this trend (Fig. S2), the observation which was an indication of the bias toward mutation-types.

### 3 Tables

Table S1: Eliminated mutations from the original PON-tstab

| | Corrected | Removed | Total |
|---|---|---|---|
| Mismatch | 48 | 4 | 52 |
| Mismatch, Duplicated | 2 | 0 | 2 |
| Duplicated | 55 | 10 | 65 |
| Triplicated | 3 | 2 | 5 |
| PDB Related (NoID) | 47 | 21 | 68 |
| PDB Related (Quality) | 0 | 14 | 14 |
| PDB Related (NoID), Mismatch | 0 | 10 | 10 |
| PDB Related (NoID), Duplicated | 1 | 48 | 49 |
| PDB Related (NoID), Triplicated | 0 | 4 | 4 |
| Total | 156 | 113 | 269 |

### 4 Figures

| | *Ali* | *Aro* | *Pol* | *Cha* |
|---|---|---|---|---|
| ***Ali*** (A, I, L, V, M, P, G) | 42 | 21 | 35 | 35 |
| ***Aro*** (F, Y, W) | 21 | 6 | 15 | 15 |
| ***Pol*** (S, T, C, N, Q) | 35 | 15 | 20 | 25 |
| ***Cha*** (D, E, H, K, R) | 35 | 15 | 25 | 20 |

Figure S1: Absolute frequencies of mutation groups based on 4L alphabet for the theoretical set of 380 mutations encompassing all possible substitutions.

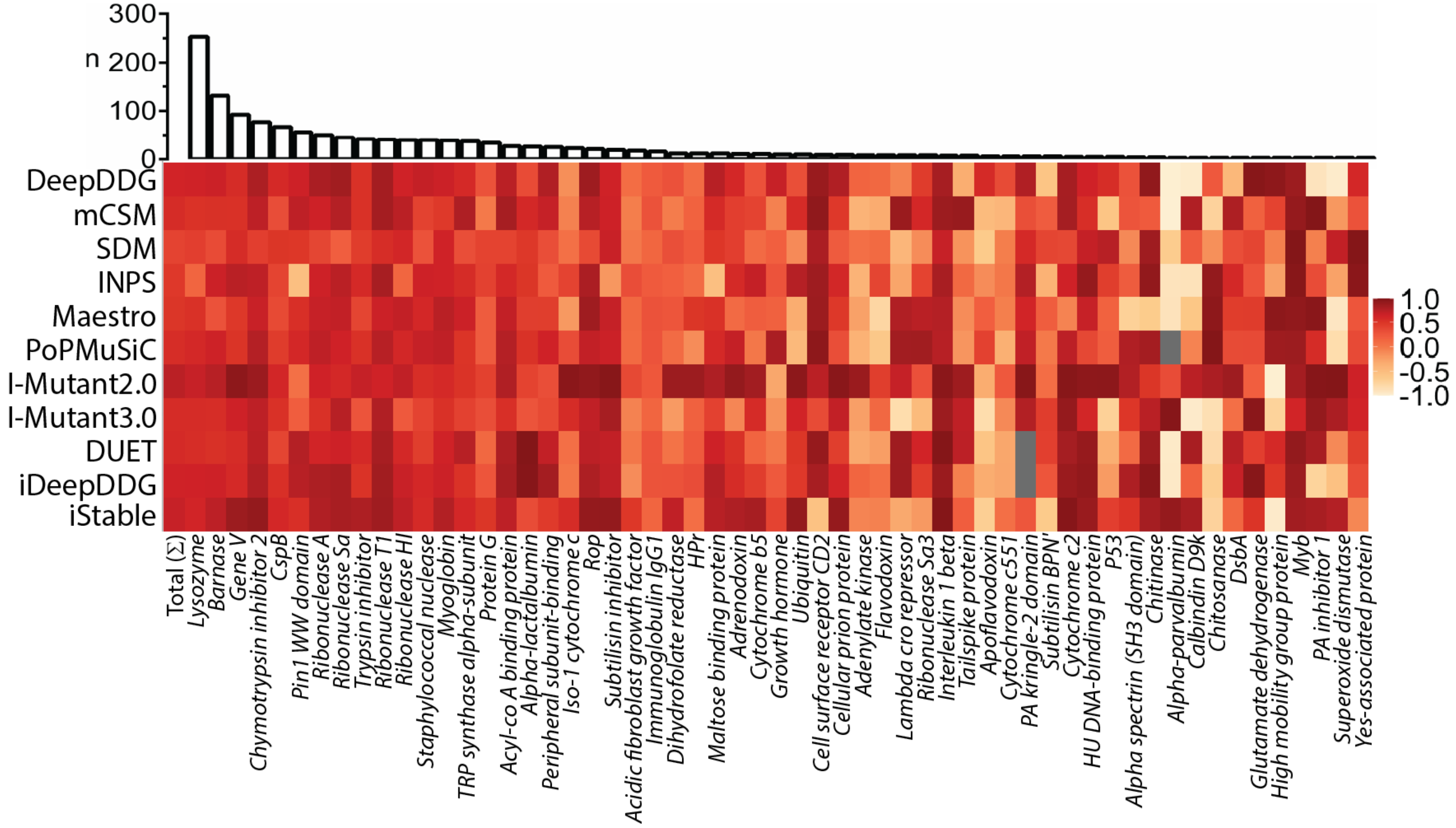


Figure S2: Scoring performance (R) based on proteins. Among 89 proteins in the curated dataset, 57 different proteins that had at least two mutations were ranked according to the number of mutations (n) in the dataset.

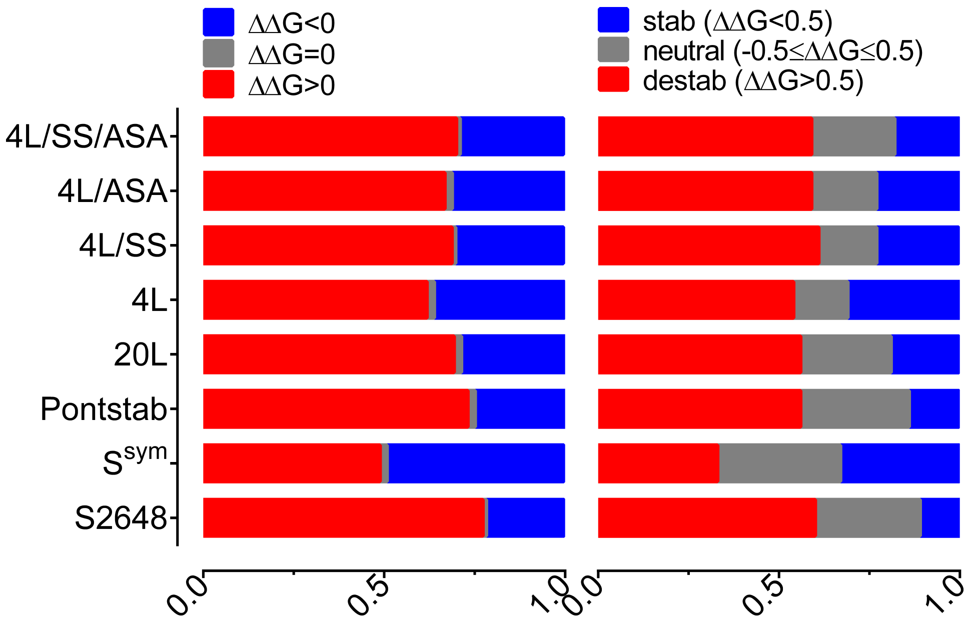


Figure S3: Relative frequencies of the stabilizing, neutral and destabilizing mutations within the datasets are displayed

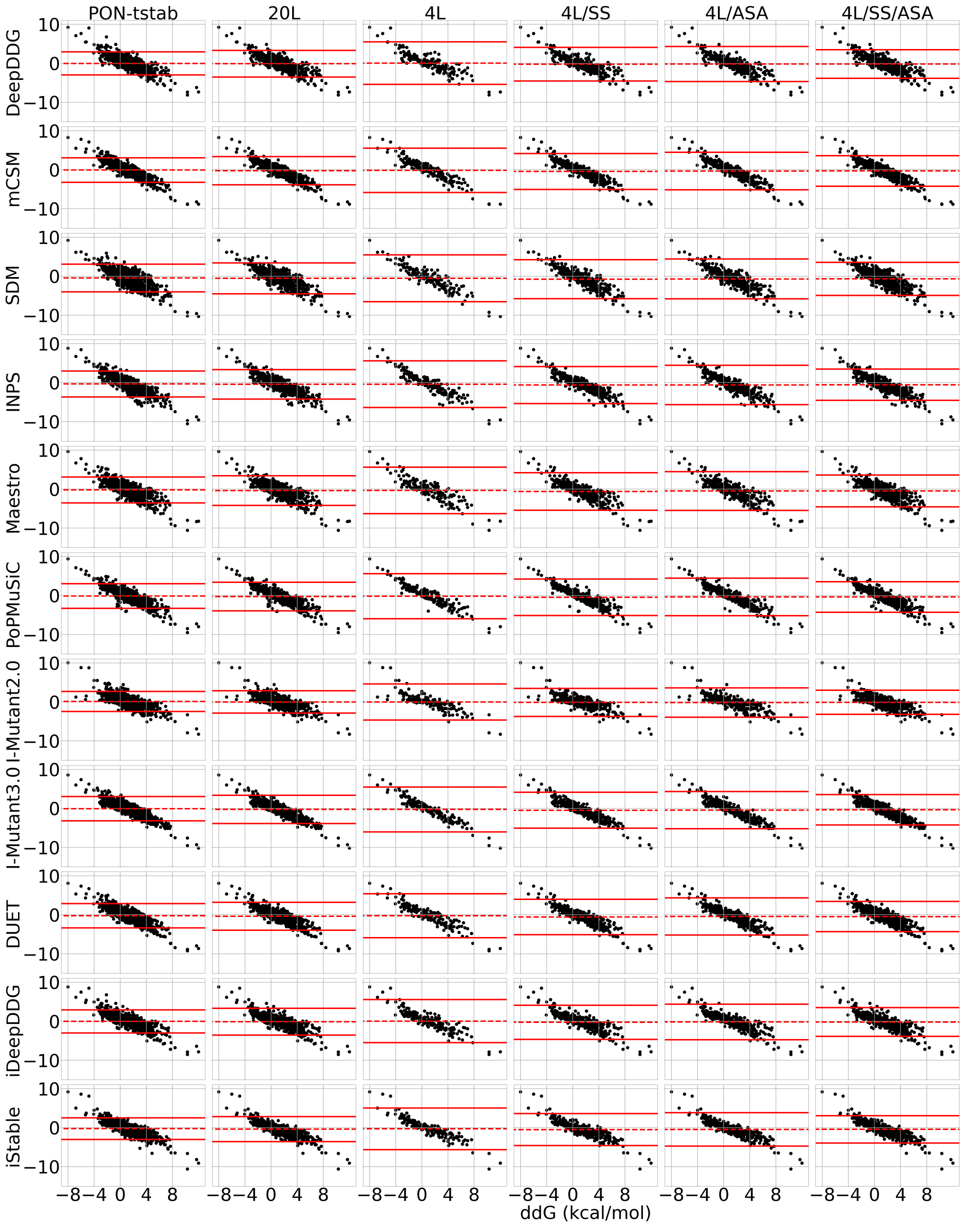


Figure S4: Bland-Altman plots of the predictors for the curated and undersampled datasets. Y axis shows the difference between predicted and experimental $\Delta\Delta G$ while the X axis represents experimental $\Delta\Delta G$. Bias (mean difference) and limits of agreement (LOA) were indicated by dashed and solid lines respectively.

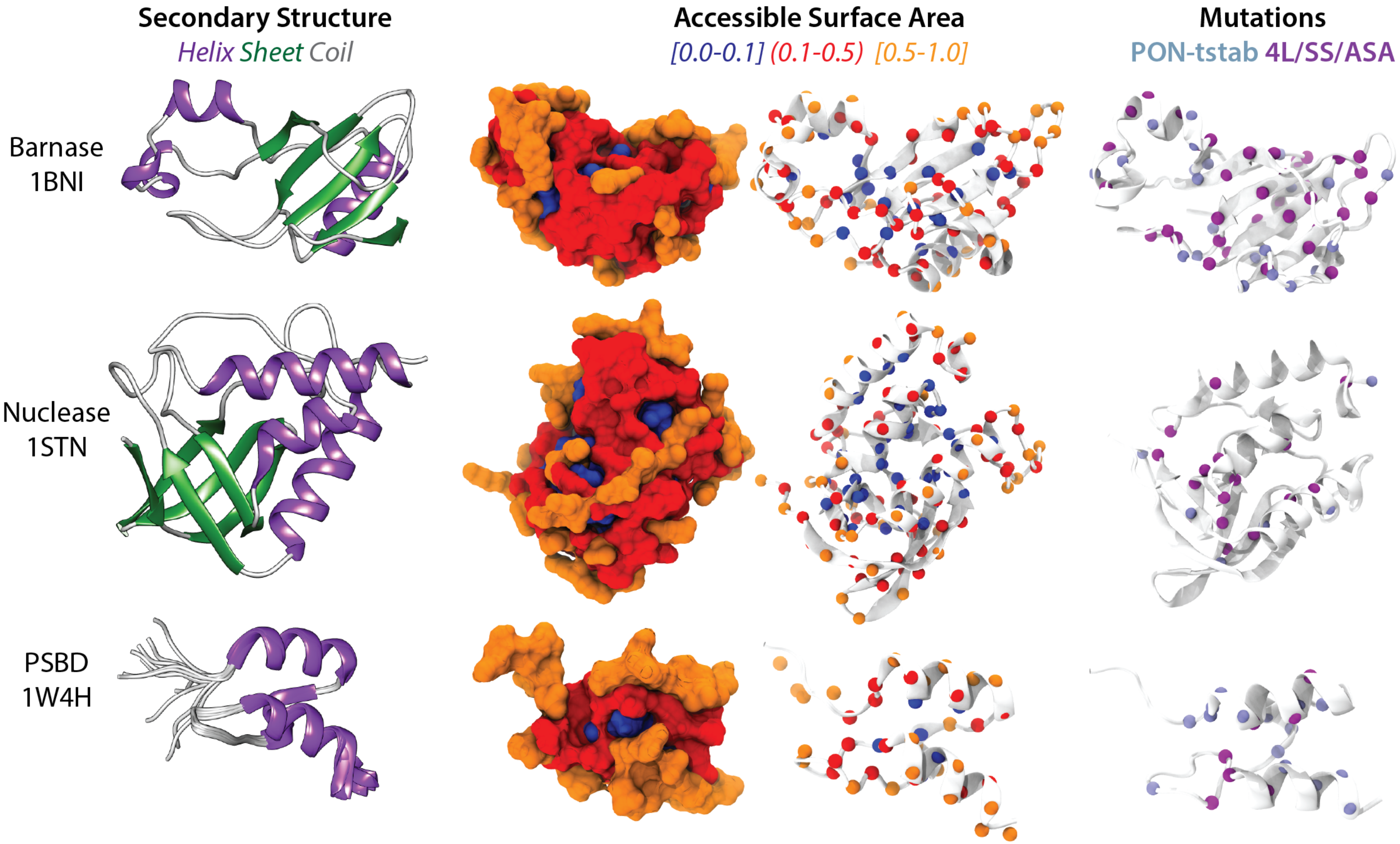


Figure S5: Structural constraints were illustrated on 3 example proteins. ASA labels were rendered by both surface and C$\alpha$ trace representations. C$\alpha$ atoms of the mutations from the PON-tstab (light) and one of the under-sampled (dark) datasets were illustrated.

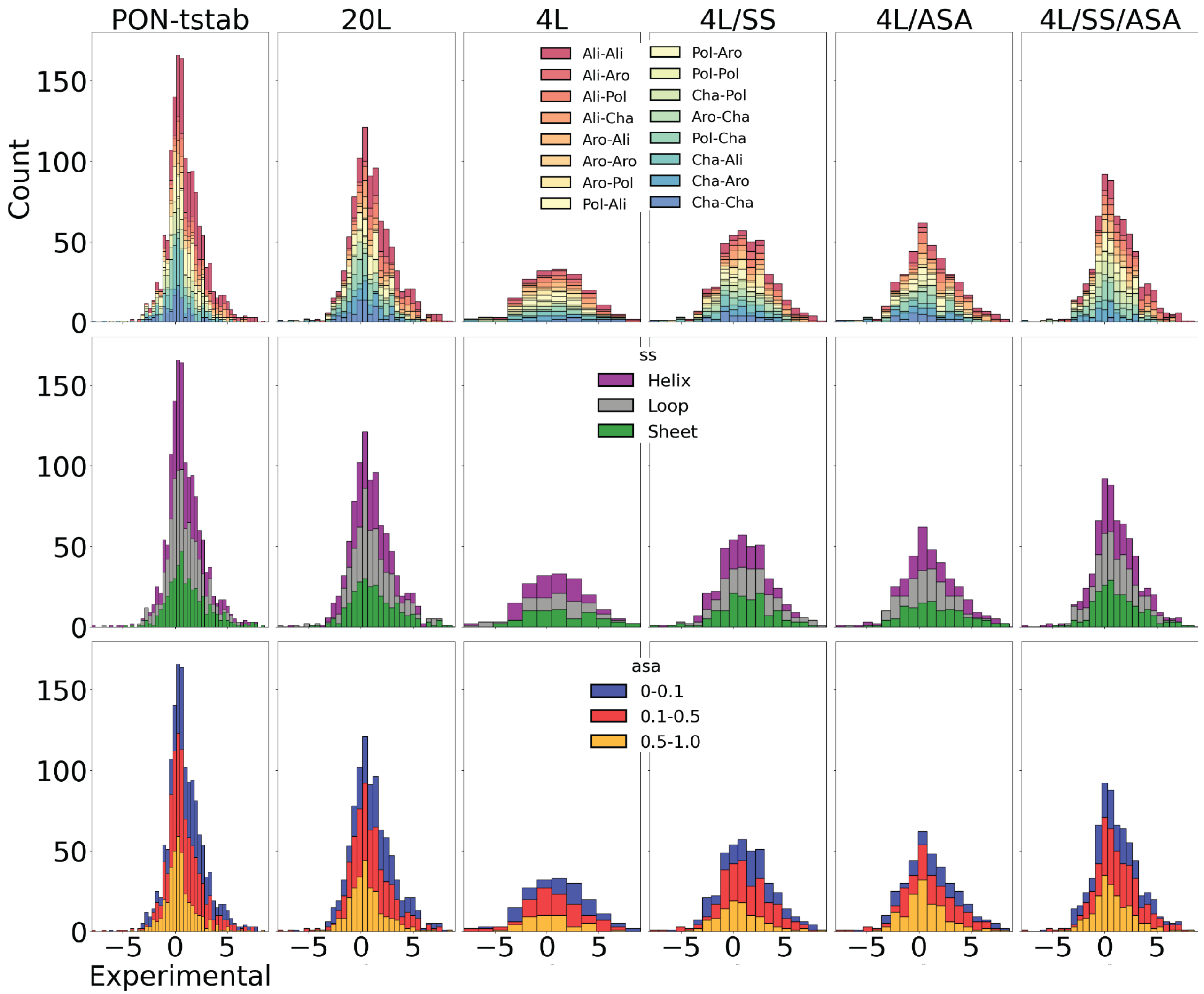


Figure S6: Scoring performance (R) based on proteins. Among 89 proteins in the curated dataset, 57 different proteins that had at least two mutations were ranked according to the number of mutations (n) in the dataset.

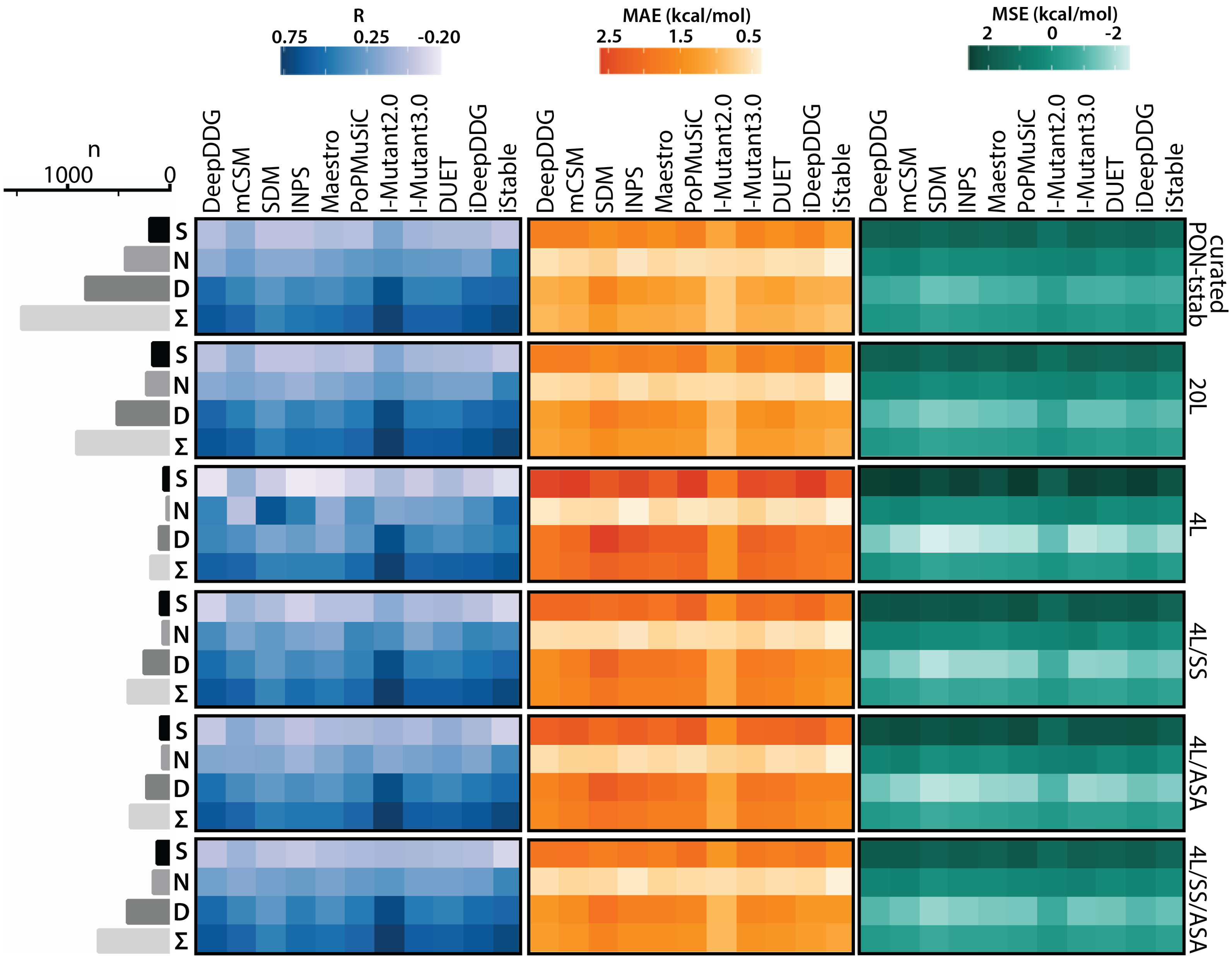


Figure S7: Performance analysis of 11 predictors on the curated PON-tstab and 5 under-sampled subsets by means of Pearson correlation coefficient (R), Mean Absolute Error (MAE) and Mean Sign Error (MSE).
